# Increasing Chlorophyll *a* Amid Stable Nutrient Concentrations in Rhode Island Lakes and Reservoirs

**DOI:** 10.1101/2020.02.11.944280

**Authors:** J. W. Hollister, D. Q. Kellogg, B. J. Kreakie, S. Shivers, W. B. Milstead, E. Herron, L. Green, A. Gold

## Abstract

Addressing anthropogenic impacts on aquatic ecosystems is a focus of lake management. Controlling phosphorus and nitrogen can mitigate these impacts, but determining management effectiveness requires long-term datasets. Recent analysis of the LAke multi-scaled GeOSpatial and temporal database for the Northeast (LAGOSNE) United States found stable water quality in the northeastern and midwestern United States, however, sub-regional trends may be obscured. We analyze a sub-regional (i.e., 3000 km^2^) trend with the University of Rhode Island’s Watershed Watch Volunteer Monitoring Program (URIWW) dataset. URIWW has collected water quality data on Rhode Island lakes and reservoirs for over 25 years. The LAGOSNE and URIWW datasets allow for comparison of water quality trends at regional and sub-regional extents, respectively. We assess regional (LAGOSNE) and state (URIWW) trends with yearly mean anomalies calculated on a per-station basis. Sub-regionally, temperature and chlorophyll *a* increased from 1993 to 2016. Total nitrogen shows a weak increase driven by low years in the early 1990s. Total phosphorus and the nitrogen:phosphorus ratio (N:P) were stable. At the regional scale, the LAGOSNE dataset shows similar trends to prior studies of the LAGOSNE with chlorophyll *a*, total nitrogen, total phosphorus, and N:P all stable over time. In short, algal biomass, as measured by chlorophyll *a* in Rhode Island lakes and reservoirs is increasing, despite stability in total nitrogen, total phosphorus, and the nitrogen to phosphorus ratio. This analysis suggests an association between lake temperature and primary production. Additionally, we demonstrate both the value of long-term monitoring programs, like URIWW, for identifying trends in environmental condition, and the utility of site-specific anomalies for analyzing for long-term water quality trends.

## 1 Introduction

Aquatic ecosystems have been altered as the result of human activities modifying nutrient cycling on a global scale (Vitousek et al. 1997, Filippelli 2008, Finlay et al. 2013). Because of their position in the landscape, lakes can function as integrators and sentinels for these anthropogenic effects (Williamson et al. 2008, Schindler 2009). Increasing nutrient inputs, particularly of nitrogen (N) and phosphorus (P), derived from intensive agriculture and densely populated urban areas have contributed to the eutrophication of many lakes (Carpenter et al. 1998, Smith 2003). This eutrophication often leads to an increase in the frequency and severity of harmful algal blooms, greater risks for human and animal health, and potential economic costs associated with eutrophic waters (Dodds et al. 2008, Paerl and Huisman 2009, Kosten et al. 2012, Michalak et al. 2013, Taranu et al. 2015, Brooks et al. 2016). To address these problems, management strategies have historically focused on reducing P inputs to lakes, but research also suggests that reducing N inputs may be more effective in certain situations (Schindler et al. 2008, Paerl et al. 2016). These studies indicate that relationships between N, P, and chlorophyll *a* exist and these relationships are spatially and temporally complex. Thus, long-term data are needed to identify trends at local, regional, and national scales.

Lake datasets that cover longer time periods and broader spatial scales are now becoming available. Programs such as the US Environmental Protection Agency’s National Lakes Assessment (NLA) provide data that allow for continental-scale water quality analysis. These data allow for analyses that can be useful for managing water resources by developing water quality criteria for N, P, and chlorophyll *a* (Herlihy et al. 2013, Yuan et al. 2014). Studying temporal trends across large spatial scales can illustrate the effects of eutrophication such as the degradation of oligotrophic systems as P increases (Stoddard et al. 2016). Broad-scale data can also be used for water quality modeling across a range of spatial scales including for predicting lake trophic state, which is indicative of ecosystem condition (Hollister et al. 2016, Nojavan et al. 2019). These trophic state models indicate that landscape variables (e.g., ecoregion, elevation, and latitude) are important and that regional trends exist. Lake-specific drivers have also been shown to be important for predicting continental-scale water quality which adds an additional layer of complexity (Read et al. 2015). Despite these challenges, it is important to study lakes at multiple spatial scales because emergent trends on regional or continental scales may or may not be present in individual lakes (Cheruvelil et al. 2013, Lottig et al. 2014).

Previous studies using regional data from the northeastern and midwestern United States (US) have investigated spatial and temporal water quality trends and have shown differences based on scale. Macro-scale (i.e., subcontinental) drivers of water quality trends are complex and may vary temporally (Lottig et al. 2017). This complexity can cause nutrient (N and P) trends to have different drivers than ratios of the nutrients (Collins et al. 2017). On a regional scale, trends of N, P, and chlorophyll *a* differ as factors such as land use and climate vary between regions, particularly when comparing the northeastern and midwestern US (Filstrup et al. 2014, 2018). Thus, it was surprising when little change in nutrients and chlorophyll *a* was reported over a 25 year period for these regions (Oliver et al. 2017). Given what is known about long-term trends in water quality within the broader region of the northeastern United States (US), we were curious if the lack of trends was also present in water quality at a sub-regional scale, using data on the 3,000 km^2^ area that encompasses a number of Rhode Island lakes and reservoirs.

Examining long-term trends in Rhode Island lakes is possible because of the data gathered by University of Rhode Island’s Watershed Watch (URIWW). URIWW is a scientist-led citizen science program founded in the late 1980s that has built a robust collaboration between URI scientists and a vast network of volunteer monitors. Volunteer monitors are trained and then collect *in situ* data as well as whole water samples during the growing season (e.g., May through October). The entire effort follows rigorous quality control/quality assurance protocols. These types of citizen science efforts allow for the collection of reliable data that in turn lead to crucial and frequently unexpected insights (Dickinson et al. 2012, Kosmala et al. 2016, Oliver et al. 2017). URIWW data contributed to the larger regional study by Oliver et al. (2017), and, also allowed us to examine the long-term trends specifically in Rhode Island.

The goals of this study were to examine ∼25 years of lake and reservoir data in Rhode Island and answer two questions. First, are there state-wide trends in total nitrogen (TN), total phosphorus (TP), total nitrogen to total phosphorus ratio (TN:TP), chlorophyll *a*, and lake temperature? Second, are water quality trends in Rhode Island similar to regional trends in the northeastern United states? Another objective of this paper was to apply existing methods for examining long-term climate records (e.g., Jones and Hulme 1996) to water quality data in order to examine long-term trends. We conducted this analysis using open data from the URI Watershed Watch program and the LAke multi-scaled GeOSpatial and temporal database for the Northeast (LAGOSNE) project and the analysis in its entirety is available for independent reproduction at https://github.com/usepa/ri_wq_trends and is archived at https://doi.org/10.5281/zenodo.3662828 (Soranno et al. 2017, Stachelek and Oliver 2017, Hollister et al. 2019).

## 2 Methods

For this study, we combined a long-term dataset on water quality of lakes in Rhode Island with a trend analysis based on water quality anomalies (i.e., measured values with the long term mean subtracted) to find increasing or decreasing annual water quality trends. Details are outlined below.

### 2.1 Study Area and Data

The study area for this analysis includes lakes and reservoirs in the state of Rhode Island where data were collected by the University of Rhode Island’s Watershed Watch program (Figure 1). The URIWW program began in 1988, monitoring 14 lakes and has now grown to include over 250 monitoring sites on over 120 waterbodies, including rivers/streams, and estuaries, with more than 400 trained volunteers. URIWW now provides more than 90% of Rhode Island’s lake baseline data and is an integral part of the state’s environmental data collection strategy. Data quality assurance and control is treated with paramount importance; volunteers are trained both in the classroom and the field, regular quality checks occur, and volunteers are provided with all the necessary equipment and supplies, along with scheduled collection dates. For freshwater lakes and reservoirs, weekly secchi depth and water temperature are recorded, along with bi-weekly chlorophyll *a* and in deep lakes (greater than 5 meters) dissolved oxygen. Water samples are collected three times per season (May through October) to be analyzed for nutrients and bacteria.

**Figure 1:**
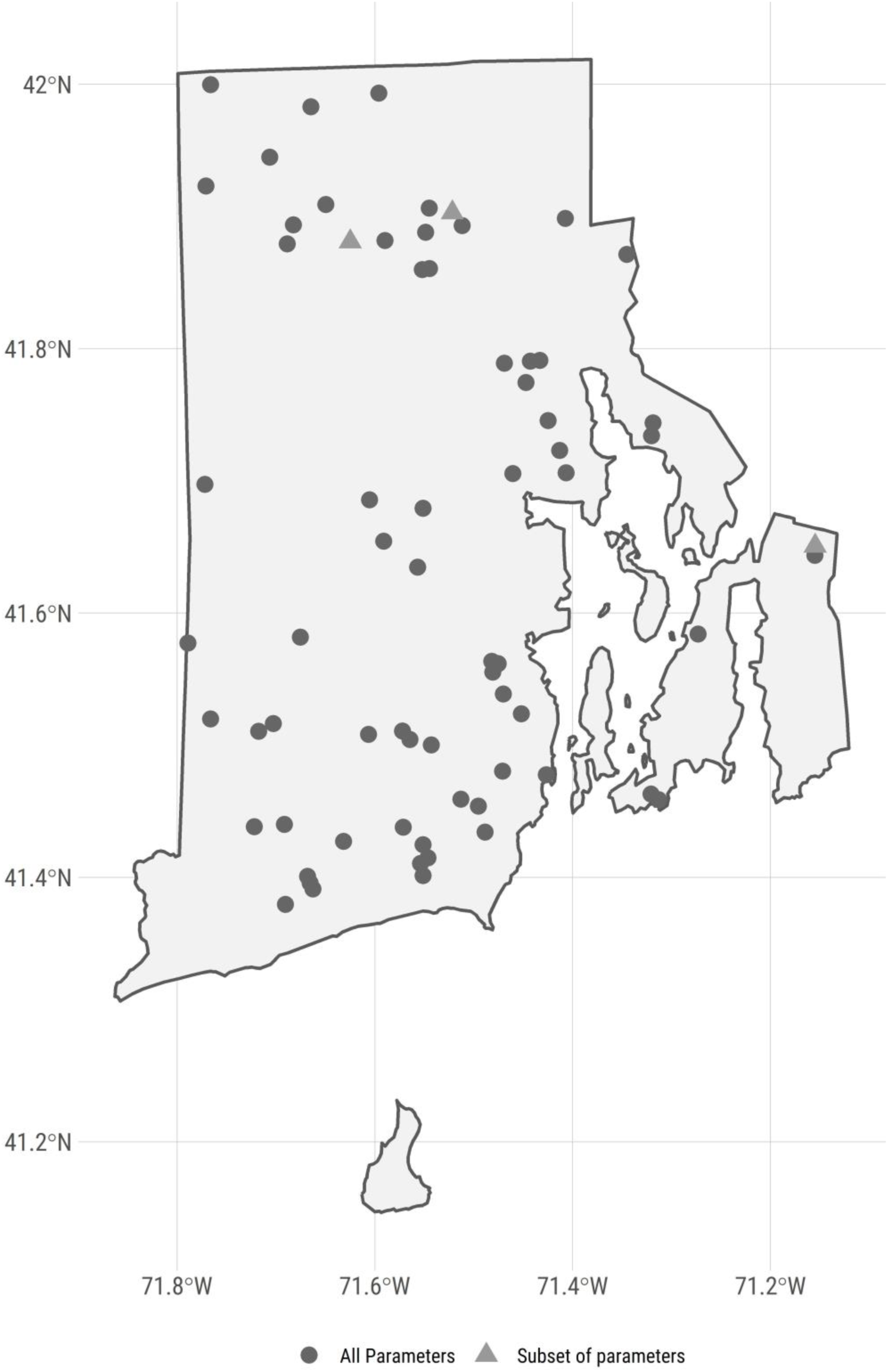
Map of URI Watershed Watch lake and reservoir sampling sites.

For this analysis, we were interested in trends in lake temperature, TN, TP, TN:TP, and chlorophyll *a*. In particular, we selected URIWW data that matched the following criteria: 1) were sampled between 1993 and 2016, 2) were sampled in May to October, 3) and were sampled at a depth of 2 meters or less. As not all sites have data for all selected years, we further filtered the data to select sites that had at least 10 years of data for a given parameter within the 1993 to 2016 time frame. The final dataset used in our analysis included 69 lakes and reservoirs. Of these sites, our filtered dataset had approximately 67 sites measured for temperature, 67 sites measured for chlorophyll *a*, 69 sites measured for TN, and 69 sites measured for TP. Of the 69 sampling sites, 66 had data for all 5 parameters. The N:P ratio was calculated by dividing the mass concentrations of total nitrogen and total phosphorus and then converting to a molar ratio by multiplying by 2.21 (e.g., atomic weight of P 30.974/atomic weight of N 14.007).

Field and analytical methods are detailed on the URIWW website at https://web.uri.edu/watershedwatch/uri-watershed-watch-monitoring-manuals/ and https://web.uri.edu/watershedwatch/uri-watershed-watch-quality-assurance-project-plans-qapps/, respectively. These methods, approved by both the state of Rhode Island and the US Environmental Protection Agency, have remained fairly consistent, although over the nearly 30 years changes did occur. When new methods were introduced, comparisons between old and new methods were conducted and in all cases no statistically significant differences were found with the new methods. Furthermore, the new methods did at times improve the limits of detection; however, this impacted a very small number (less than 1%) of measurements in this study. We did run our analyses (see **Water Quality Trend Analysis** section) with all data and with only those data greater than the detection limit. There was no change in the trend analysis and thus, the results we report are for all data as originally reported in the URIWW dataset. Given these results, we assume the data to be consistent across the reported time period and appropriate for a long term assessment of trends.

Prior studies have modeled water quality trends across a larger region of the northeastern US that included 17 states including Minnesota, Wisconsin, Iowa, Missouri, Illinois, Indiana, Michigan, Ohio, Pennsylvania, New York, New Jersey, Connecticut, Massachusetts, Rhode Island, Vermont, New Hampshire, and Maine (Soranno et al. 2015, Oliver et al. 2017). We repeated our analysis (see **Water Quality Trend Analysis** section) with the same dataset used by Oliver et al. (2017), the LAGOSNE dataset (Soranno et al. 2015, 2017, Stachelek and Oliver 2017). Temperature data were not available, thus we examined trends, using our analytical methods, for TN, TP, TN:TP, and chlorophyll *a* from the LAGOSNE dataset. We used the same selction criteria on the LAGOSNE dataset as was applied to the URIWW data.

### 2.2 Water Quality Trend Analysis

There are many different methods for analyzing time series data for trends. Environmental data are notoriously “noisy” and one of the difficulties that is encountered with multiple sampling locations is how to identify a trend while there is variation within a sampling site as well as variation introduced by differing start years for sampling among the many sites. For instance, if long-term data on water quality were collected more frequently in early years from more pristine waterbodies, then a simple comparison of raw values over time might show a decrease in water quality, which could be misleading if later sampling occurred on both pristine and more eutrophic water bodies. Thus, it is necessary to account for this type of within-site and among-site variation, using methods similar to those used to analyze long-term temperature trends using temperature anomalies (e.g., Jones and Hulme 1996). The general approach we used calculates site-specific deviations from a long-term mean over a pre-determined reference period. This allowed all sites to be shifted to a common baseline and the deviations, or anomalies, indicate change over the specified reference period. We refer to this method as “site-specific anomalies”.

#### 2.2.1 Summarizing site-specific anomalies

Methods for calculating the site-specific anomalies and the yearly means are as follows and are presented graphically in Figure 2. Additionally, an example R script, `schematic_anomaly.R` and example dataset, `schematic.csv` to recreate and demonstrate the calculations in Figure 2 is available from at https://github.com/usepa/ri_wq_trends and is archived at https://doi.org/10.5281/zenodo.3662828 (Hollister et al. 2019).

**Figure 2:**
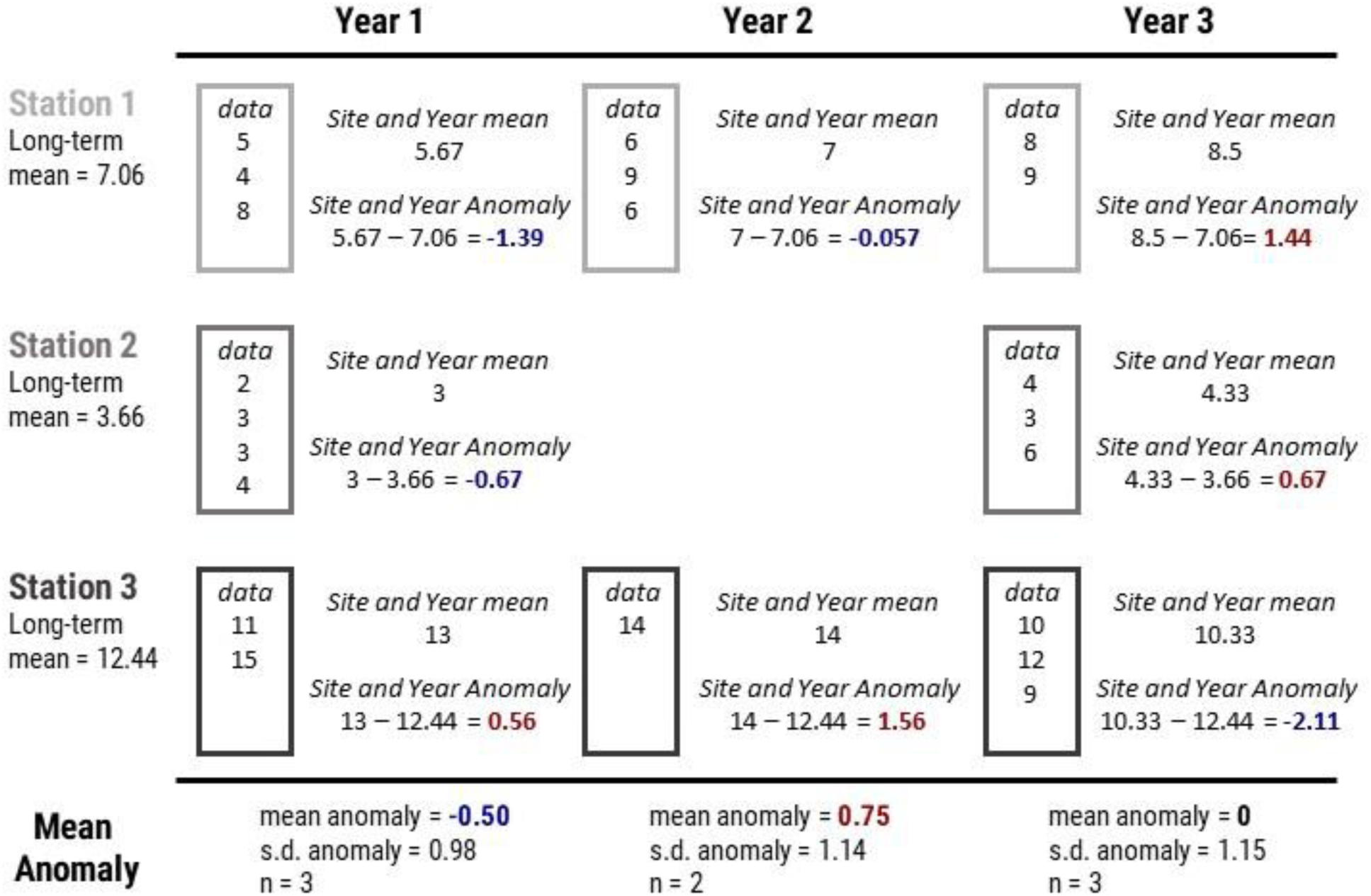
Example calculation of the site-specific anomalies and yearly mean anomalies.

The general steps, outlined in Figure 2 and listed below, are repeated for each of the water quality parameters.

1. For each site, calculate the annual means, producing a single mean value for each site and year. This step prevents bias from pseudoreplication of multiple measurements of the same site in a given year (Hurlbert 1984). The per site means across years are assumed to be independent.
2. Calculate the long-term reference mean for each site. This results in a single long-term mean for each of the sites.
3. Calculate the anomaly for each annual mean at each site by subtracting the annual and reference means.
4. Summarize by calculating the mean anomaly per year for the entire group of sites. The resultant values are analyzed for a trend over time.

#### 2.2.2 Linear regression on annual mean anomalies

Testing for a regression slope being different than zero can be used to test for monotonic trends in water quality data (Helsel and Hirsch 2002). We used these standard procedures to test for positive or negative trends in lake temperature, chlorophyll *a*, TN, TP and TN:TP. For each parameter, we fit a regression line to the anomalies as a function of year and tested the null hypothesis that no trend existed (e.g., *β*_1_ = 0). The slope of this line provides information on the mean yearly change of that paramter over the time period studied. Traditionally, trends would be determined by assessing “significance” but recent guidelines suggest not using arbitrary p-value cut-offs to assesses significance (Wasserstein et al. 2016). Our interpretation of the trends attempts to follow this advice and we assess trends with the information provided by the magnitude of the slopes, the p-values, and our understanding of the processes involved.

#### 2.2.3 Comparison of Rhode Island to the region

Oliver et al. (2017) used hierarchical linear models and showed relatively stable water quality in the lakes of the northeastern United States. While the University of Rhode Island’s Watershed Watch data were included in this regional study, we hypothesized that in the case of Rhode Island regional trends were masking sub-regional trends. Therefore, we decided to reanalyze the LAGOSNE data to compare the trends at the regional scale to the trends at the Rhode Island state scale using the site-specific anomaly and trend analysis approach outlined above.

## 3 Results

During the period of 1993 to 2016, Rhode Island lakes and reservoirs in our dataset had a mean lake temperature of 21.9 °C, mean TN of 600 μg/l, mean TP of 24 μg/l, mean TN:TP ratio of 84.17 molar, and mean chlorophyll *a* of 10.1 μg/l (Table 1).

**Table 1:** Summary statistics for URI Watershed Watch data from 1993 to 2016.

| Parameter | Units | Mean | Median | Max | Std. Dev |
| --- | --- | --- | --- | --- | --- |
| Temperature | °C | 21.9 | 22.2 | 29 | 1.9 |
| Total Nitrogen | µg/l | 600 | 475 | 4670 | 425 |
| Total Phosphorus | µg/l | 24 | 15 | 325 | 30 |
| N:P | molar | 84.17 | 71.08 | 827.2 | 57.9 |
| Chlorophyll | µg/l | 10.1 | 4.5 | 666.2 | 22.1 |

For lakes and reservoirs in the larger region represented by the LAGOSNE States, mean TN was 855 μg/l, mean TP was 32 μg/l, mean TN:TP ratio was 90.37 molar, and mean chlorophyll *a* was 16.8 μg/l (Table 2).

**Table 2:** Summary statistics for LAGOSNE data from 1993 to 2016.

| Parameter | Units | Mean | Median | Max | Std. Dev |
| --- | --- | --- | --- | --- | --- |
| Total Nitrogen | µg/l | 855 | 560 | 16780 | 1205 |
| Total Phosphorus | µg/l | 32 | 16 | 1200 | 54 |
| N:P | molar | 90.37 | 59.18 | 88474 | 1029 |
| Chlorophyll | µg/l | 16.8 | 6.2 | 696 | 30.4 |

### 3.1 State-wide trends in water quality

Mean annual temperature anomalies in lakes and reservoirs appears to be increasing (slope = 0.053, p = 0.0062) with the majority of years with mean temperature greater than the long-term mean occurring in recent years (Figure 3). Chlorophyll *a* is also showing an increasing trend over time (slope = 0.29, p = 0.0000008) and with the exception of a slightly above-average year in 2003, the above-average years have all occurred in the most recent years (Figure 4A.).

**Figure 3:**
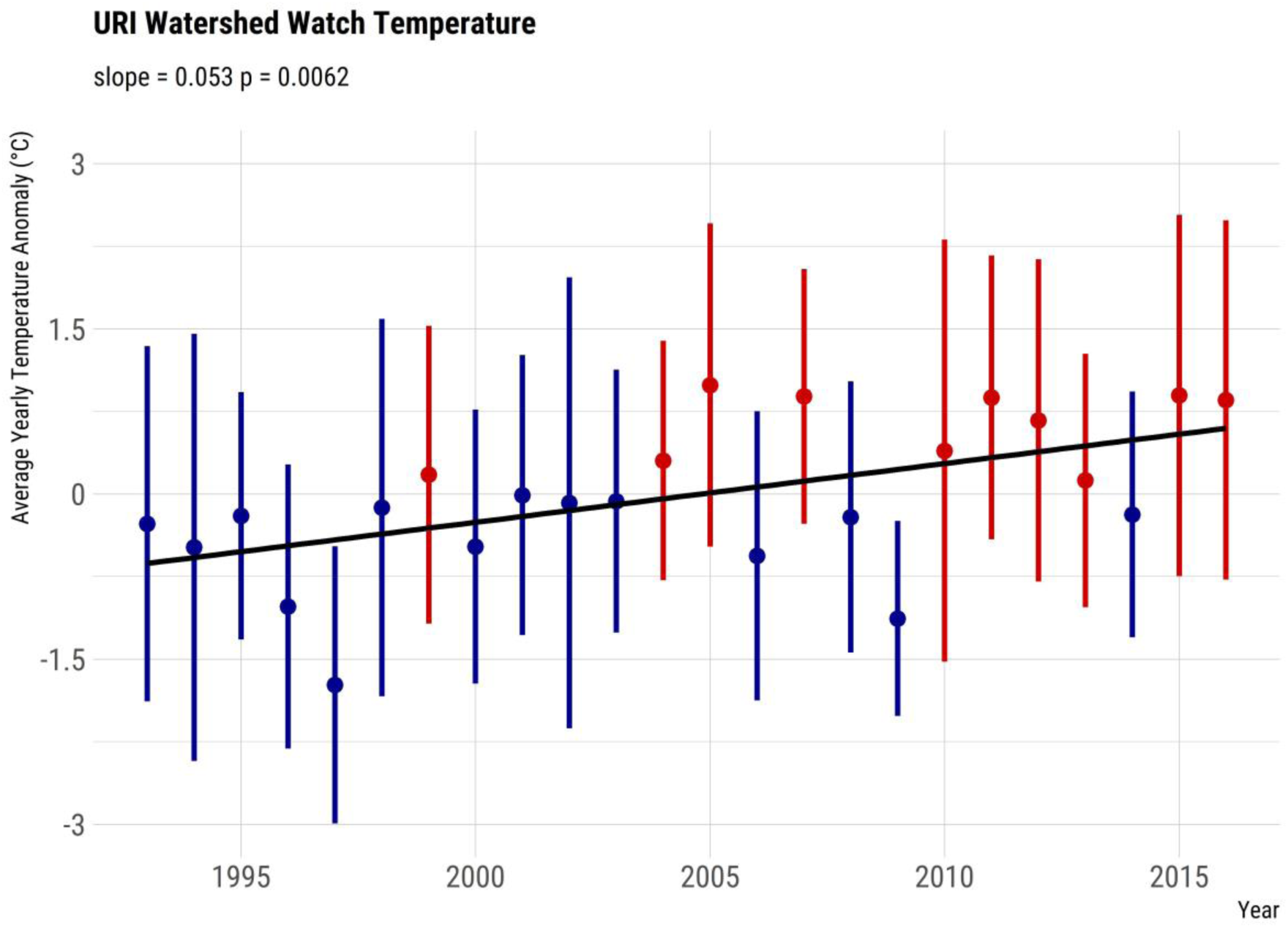
Yearly trend over 20+ years of lake temperature (mean anomaly) in Rhode Island lakes and reservoirs. Points are means of site-specific anomalies and ranges are standard deviations of site-specific anomalies. Blue indicates yearly site-specific anomalies that were, on average, below the site-specific long-term means. Red indicates yearly site-specific anomalies that were, on average, above the site-specific long-term means.

**Figure 4:**
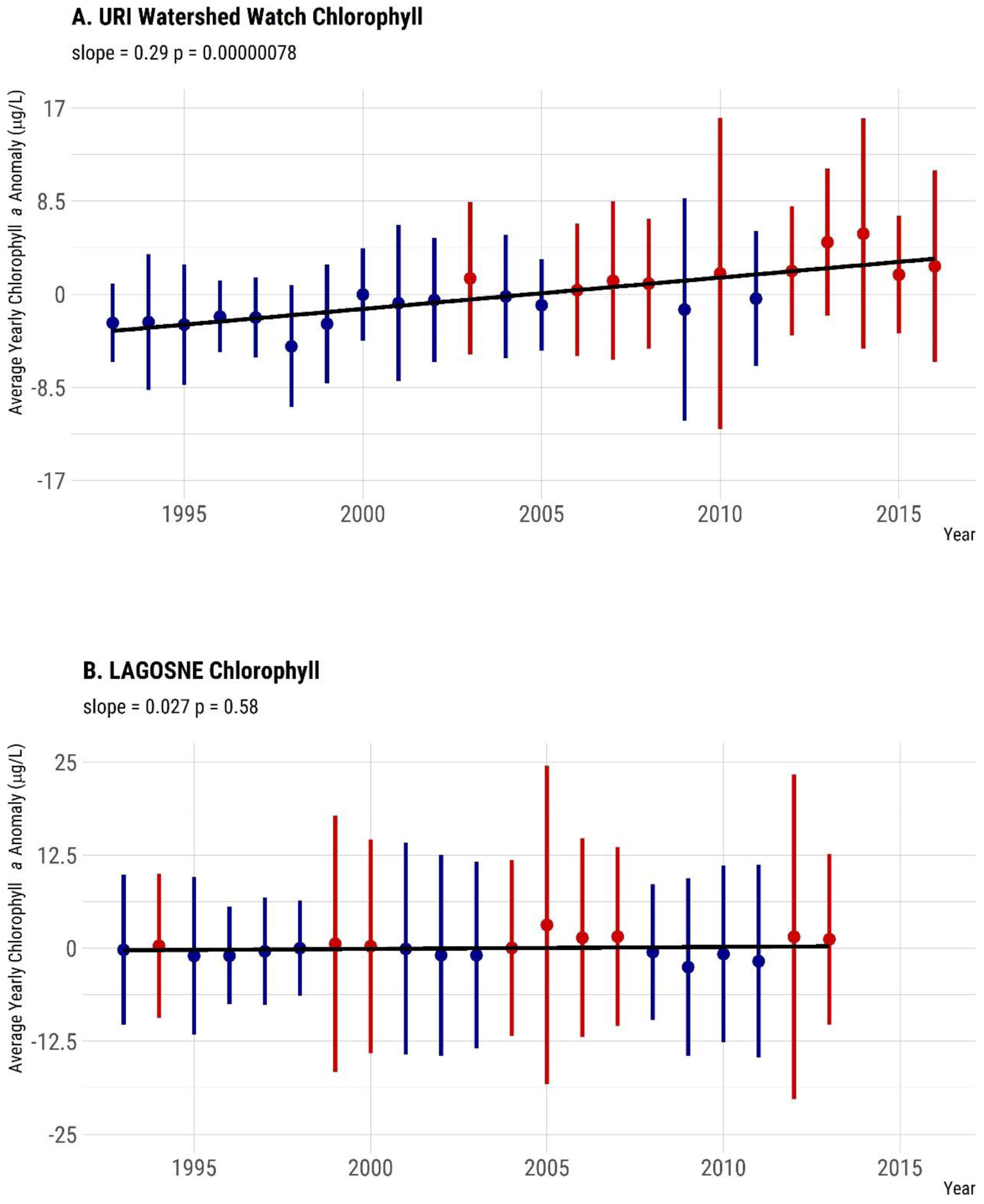
Yearly trend over 20+ years of chlorphyll a (mean anomaly). Panel A. Yearly mean chlorophyll a anomalies from the URI Watershed Watch data. Panel B. Yearly mean chlorophyll a anomalies from the LAGOSNE dataset. Points are means of site-specific anomalies and ranges are standard deviations of site-specific anomalies. Blue indicates yearly site-specific anomalies that were, on average, below the site-specific long-term means. Red indicates yearly site-specific anomalies that were, on average, above the site-specific long-term means.

Mean annual trends for nutrients were weaker or showed no trend over time. The data suggest a positive trend in TN (slope = 3.8, p = 0.00022); however, that perceived trend is driven by the lower than mean TN values in 1993 and 1994 (Figure 5A.). Since 1995, the yearly trend shows a lower increase over time (slope = 2.5, p = 0.0067). TP does not show a trend over time in the yearly anomalies (slope = 0.11, p = 0.062) and years that are over or under the mean are more evenly distributed over the years (Figure 6A.). The pattern is the same for the TN:TP ratio (slope = 0.18, p = 0.71) with little evidence suggesting a change in the concentrations of TN relative to the concentrations of TP (Figure 7A.). Data for all figures are available as a comma-separated values file, yearly_average_anomaly.csv from at https://github.com/usepa/ri_wq_trends and is archived at https://doi.org/10.5281/zenodo.3662828 (Hollister et al. 2019).

**Figure 5:**
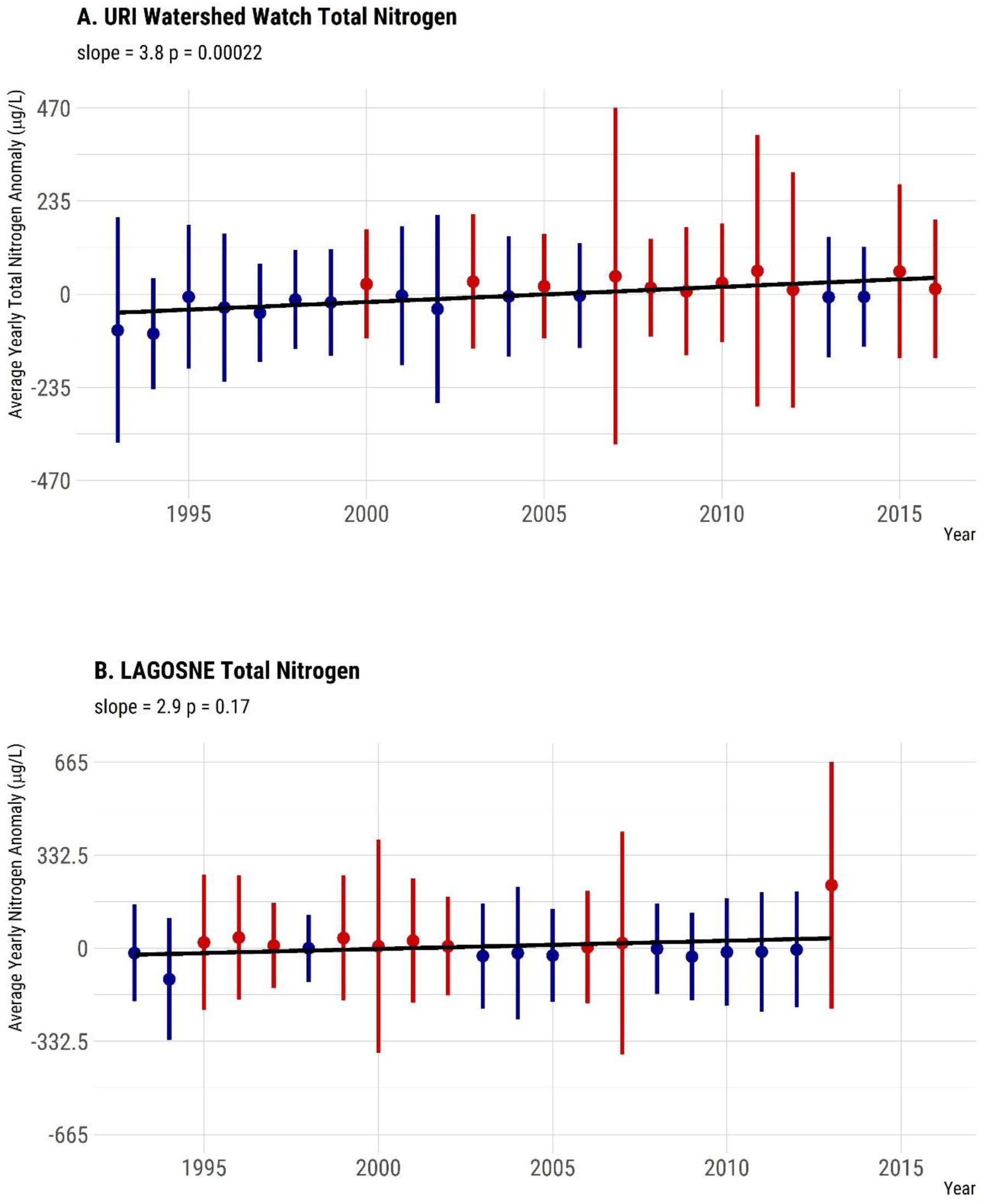
Yearly trend over 20+ years of TN (mean anomaly). Panel A. Yearly mean TN anomalies from the URI Watershed Watch dataset. Panel B. Yearly mean TN anomalies from the LAGOSNE dataset. Points are means of site-specific anomalies and ranges are standard deviations of site-specific anomalies. Blue indicates yearly site-specific anomalies that were, on average, below the site-specific long-term means. Red indicates yearly site-specific anomalies that were, on average, above the site-specific long-term means.

**Figure 6:**
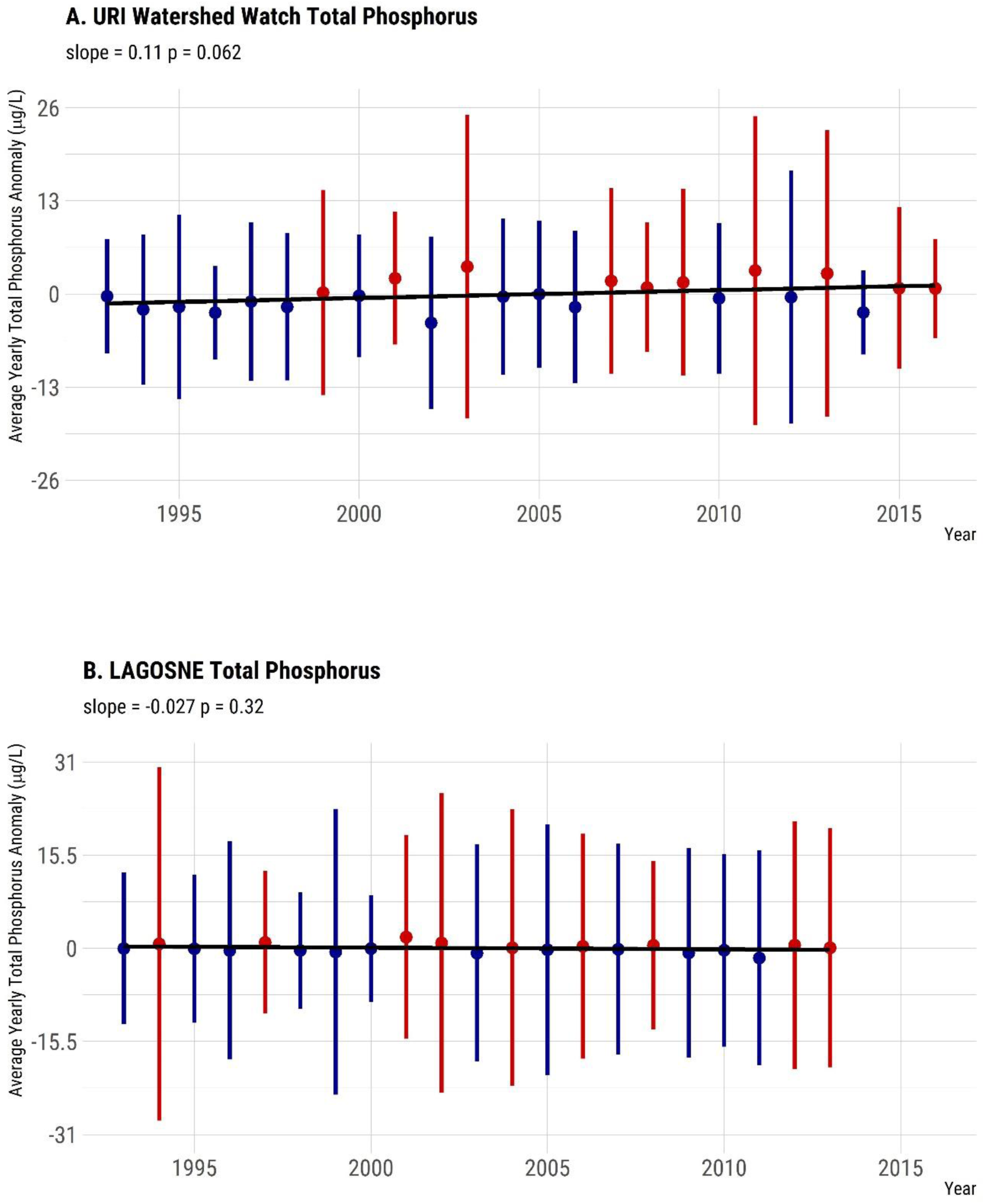
Yearly trend over 20+ years of TP (mean anomaly). Panel A. Yearly mean TP anomalies from the URI Watershed Watch dataset. Panel B. Yearly mean TP anomalies from the LAGOSNE dataset. Points are means of site-specific anomalies and ranges are standard deviations of site-specific anomalies. Blue indicates yearly site-specific anomalies that were, on average, below the site-specific long-term means. Red indicates yearly site-specific anomalies that were, on average, above the site-specific long-term means.

**Figure 7:**
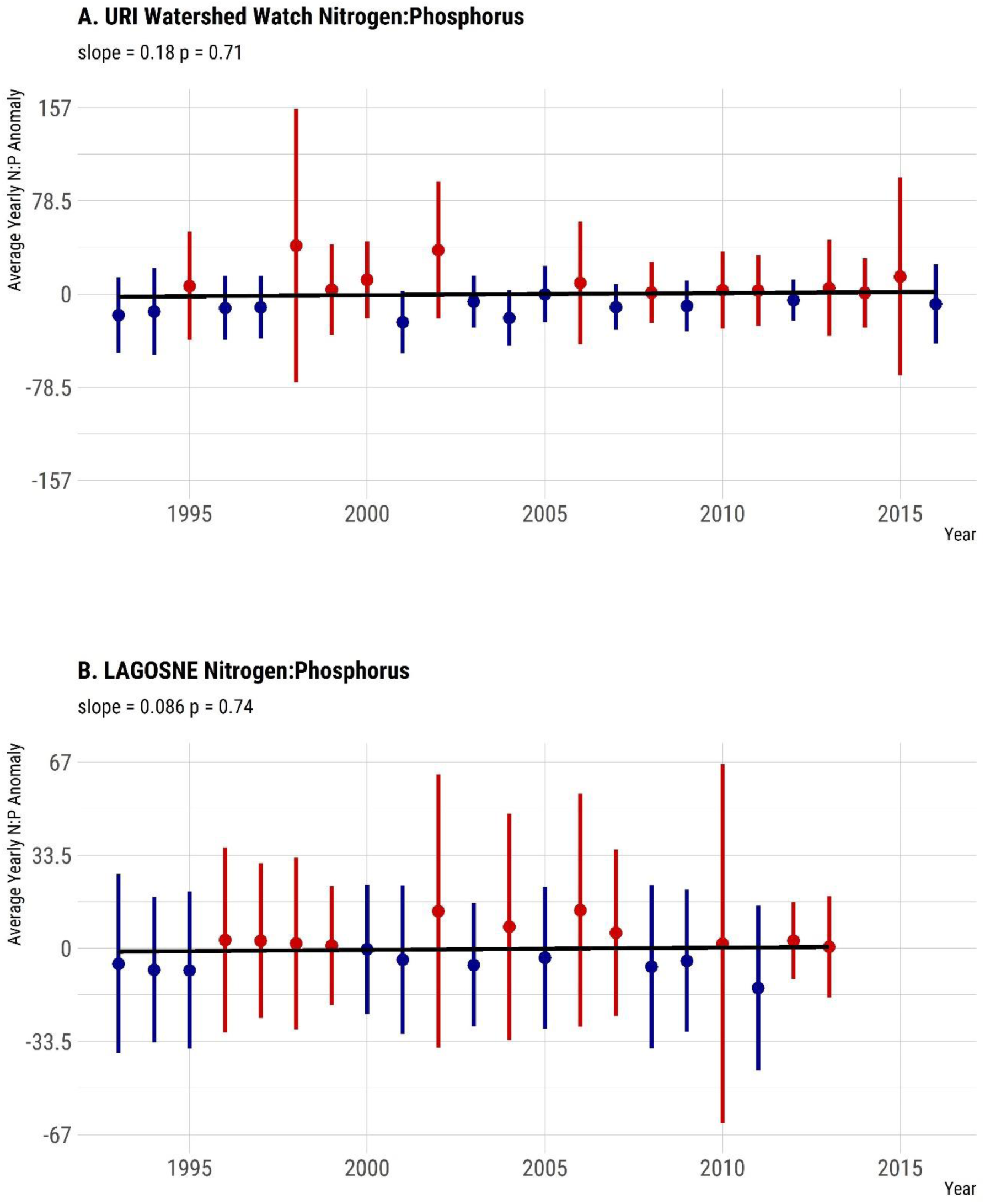
Yearly trend over 20+ years of the TN:TP ratio (mean anomaly). Panel A. Yearly mean TN:TP ratio anomalies from the URI Watershed Watch dataset. Panel B. Yearly mean TN:TP ratio anomalies from the LAGOSNE dataset. Points are means of site-specific anomalies and ranges are standard deviations of site-specific anomalies. Blue indicates yearly site-specific anomalies that were, on average, below the site-specific long-term means. Red indicates yearly site-specific anomalies that were, on average, above the site-specific long-term means.

### 3.2 Regional trends in water quality

In general, there was little evidence to suggest broad regional changes. Chlorophyll *a* showed a very weak positive trend (slope = 0.027, p = 0.58, Figure 4B.), TP showed a slight decreasing trend (slope = -0.027, p = 0.32, Figure 6B.), TN showed a slight positive trend (slope = 2.9, p = 0.17, Figure 5B.) and the TN:TP showed little change (slope = 0.086, p = 0.74, Figure 7B.)

## 4 Discussion and conclusions

Our sub-regional analysis indicates that even when nutrient regimes exhibit relative stability (i.e., neither increasing nor decreasing over time), increases in primary production, as measured by chlorophyll *a*, can occur. Over the same period we also demonstrate long-term warming of Rhode Island lakes and reservoirs. Chlorophyll has increased, on average, 0.29 μg/L per year over the 23 years of our analysis, while temperature has increased 0.053 °C per year over the same period. This suggests that the observed increase in productivity, as measured by chlorophyll *a*, may be a result of warming waters and not a response to changes in nutrient condition. Also, geographic extent does indeed matter when trying to identify long-term water quality trends. Similar to the results of Oliver et al. (2017) our analysis shows little increasing trend in chlorophyll *a* at the regional scale (e.g., northeastern and mid-western United States). However, at the local scale of the state of Rhode Island, there is a clear increasing trend in chlorophyll *a*.

### 4.1 Trends

As previously mentioned, both temperature and chlorophyll *a* show increasing trends from 1993 to 2016 in Rhode Island lakes and reservoirs; while total nutrients and the TN:TP ratio are all relatively stable. While TN showed a weak positive trend, that trend was largely driven by the unusually low years for TN in 1993 and 1994. With those removed the positive trends weakens considerably. The general picture in Rhode Island appears to be one of little to no change in phosphorus, a very weak positive trend in nitrogen and little to no change in the TN:TP ratio. Furthermore, it has been shown that productivity in freshwater systems is likely a function of both phosphorus and nitrogen (Paerl et al. 2016). Thus, the increasing chlorophyll *a* in the face of stable TN:TP ratio suggests that the increase is being driven by something other than nutrients. We interpret these results as relative stability in nutrients in Rhode Island lakes and reservoirs.

Stable nutrient regimes may be partly explained by efforts to curb nutrient loadings, for example through voluntary and state wide mandatory bans on phosphates in laundry detergent which were implemented in Rhode Island in 1995 (Rhode Island State Legislature 1995, Litke 1999). However, in many lakes there are still likely sufficient nutrients present to allow for increases in chlorophyll *a*. Additionally, these results point to the fact that chlorophyll *a* and algal biomass is driven by processes operating at different scales. For instance, nutrient management is largely a local to watershed scale effort, but may also be regional as atmospheric nitrogen deposition can be a significant source of nitrogen (Boyer et al. 2002). Similarly, warming lakes are driven by broader climate patterns, yet waterbody-specific factors such as the percent of a catchment that is impervious surface and lake morphology can also impact temperature (Nelson and Palmer 2007). In short, differences in regional and state level trends are driven by complex and multi-scale processes.

In addition to the annualized trends of the five variables we address with this study, there are other trends that may be of interest. For example, trends for water quality at finer temporal scales such as monthly or seasonal trends may be different than the annual trends we analyzed. Anecdotal evidence in Rhode Island points to warmer temperature earlier and later in the year and suggests a lengthening of the growing season. Furthermore, preliminary analysis of the URIWW data back this up with mean temperature for May 1993 to May 1995 cooler by nearly a degree than mean temperature for May 2014 through May 2016. Additionally, it may be possible that the current trophic state of a given waterbody may partly explain the chlorophyll *a* changes in that lake. For instance, are oligotrophic lakes showing stronger trends than eutrophic lakes or are all lakes showing similar trends regardless of current trophic status? Lastly, changes in rainfall, extreme weather events, or other climate mediated factors can also be playing a role in increasing chlorophyll in Rhode Island lakes and reservoirs. These questions are beyond the scope of this study, but all warrant further, careful investigation.

### 4.2 Management implications

There are several broader management implications from the results of our analysis and of examining long-term water quality trends in general. In particular, this analysis provides much needed information about the long-term effects of current nutrient control efforts at lake-specific and sub-regional scales and identifies areas where additional information is required or a change in management approaches may be needed. First, as more long-term datasets become available, it is important for managers and stakeholders to receive feedback on long-term water quality trends at multiple spatial scales. Specifically for this study, the results provide feedback to long time volunteer monitors, highlighting the importance of volunteer monitoring programs. Second, with information on long-term trends, it is possible to adapt management approaches to address areas of concern. Our results show increasing chlorophyll *a* even though the general long-term nutrient trends have been stable. This suggests the need to further reduce nutrients to compensate for warmer water temperatures and possible longer growing seasons.

There are several possible approaches to further reduce nutrient loads (Yang and Lusk 2018). First, nutrient load reductions may be possible through source controls and enhanced entrainment and treatment of ground and surface waters transporting nutrients to receiving waters (Kellogg et al. 2010). Green infrastructure approaches are one way to possibly achieve both goals (Pennino et al. 2016, Reisinger et al. 2019). Additionally, there is potential for within-lake approaches such as the restoration of freshwater mussels to waterbodies that historically had those species. Some studies using freshwater mussels have shown reductions in both nutrients and algal biomass (Kreeger et al. 2018).

### 4.3 Data analysis approach

The analysis approach we used here, site-specific anomalies, is not a novel method and does have a long history in the analysis of trends in climate (Jones and Hulme 1996, Jones et al. 1999, Hansen et al. 2006, 2010). However, using it to examine water quality trends is a new application of the technique, as we could find little evidence of using it specifically for water quality trends. We built on these methods and adapted them for use with long-term water quality trends. While other methods are valid and robust (e.g., Oliver et al. 2017), we chose mean site-specific anomalies as they can provide readily interpretable results, especially for communicating to general audiences. For instance, reporting the changes in anomalies allows us to look at changes in the original units. With our analysis, the slope of the regression line for temperature suggests a mean yearly increase of 0.053 °C and the slope of the regression line for chlorophyll *a* shows a mean yearly increase of 0.29 μg/l. Additionally, the site-specific anomalies are robust to variations in sampling effort and in the timing of inclusion of given sampling locations (e.g., added later in a time period or removed). Lastly, this analysis is only possible because of the availability of sound, long-term data on water quality in Rhode Island. Without the URIWW data and the commitment and participation of more than 2500 volunteers over the years, our analyses would have been impossible. Going forward, it is important to appreciate the role that volunteer monitoring and citizen science programs can play in capturing and better understanding long term environmental trends.

## Notes

https://github.com/usepa/ri_wq_trends

https://doi.org/10.5281/zenodo.3662828

## Bibliography

Boyer, E. W., C. L. Goodale, N. A. Jaworski, and R. W. Howarth. 2002. Anthropogenic nitrogen sources and relationships to riverine nitrogen export in the northeastern usa. Biogeochemistry 57:137–169.

Brooks, B. W., J. M. Lazorchak, M. D. Howard, M.-V. V. Johnson, S. L. Morton, D. A. Perkins, E. D. Reavie, G. I. Scott, S. A. Smith, and J. A. Steevens. 2016. Are harmful algal blooms becoming the greatest inland water quality threat to public health and aquatic ecosystems? Environmental Toxicology and Chemistry 35:6–13.

Carpenter, S. R., N. F. Caraco, D. L. Correll, R. W. Howarth, A. N. Sharpley, and V. H. Smith. 1998. Nonpoint pollution of surface waters with phosphorus and nitrogen. Ecological Applications 8:559–568.

Cheruvelil, K., P. Soranno, K. Webster, and M. Bremigan. 2013. Multi-scaled drivers of ecosystem state: Quantifying the importance of the regional spatial scale. Ecological Applications 23:1603–1618.

Collins, S. M., S. K. Oliver, J.-F. Lapierre, E. H. Stanley, J. R. Jones, T. Wagner, and P. A. Soranno. 2017. Lake nutrient stoichiometry is less predictable than nutrient concentrations at regional and sub-continental scales. Ecological applications 27:1529–1540.

Dickinson, J. L., J. Shirk, D. Bonter, R. Bonney, R. L. Crain, J. Martin, T. Phillips, and K. Purcell. 2012. The current state of citizen science as a tool for ecological research and public engagement. Frontiers in Ecology and the Environment 10:291–297.

Dodds, W. K., W. W. Bouska, J. L. Eitzmann, T. J. Pilger, K. L. Pitts, A. J. Riley, J. T. Schloesser, and D. J. Thornbrugh. 2008. Eutrophication of us freshwaters: Analysis of potential economic damages. ACS Publications.

Filippelli, G. M. 2008. The global phosphorus cycle: Past, present, and future. Elements 4:89–95.

Filstrup, C. T., T. Wagner, S. K. Oliver, C. A. Stow, K. E. Webster, E. H. Stanley, and J. A. Downing. 2018. Evidence for regional nitrogen stress on chlorophyll a in lakes across large landscape and climate gradients. Limnology and Oceanography 63:S324–S339.

Filstrup, C. T., T. Wagner, P. A. Soranno, E. H. Stanley, C. A. Stow, K. E. Webster, and J. A. Downing. 2014. Regional variability among nonlinear chlorophyll—phosphorus relationships in lakes. Limnology and Oceanography 59:1691–1703.

Finlay, J. C., G. E. Small, and R. W. Sterner. 2013. Human influences on nitrogen removal in lakes. Science 342:247–250.

Hansen, J., R. Ruedy, M. Sato, and K. Lo. 2010. Global surface temperature change. Reviews of Geophysics 48.

Hansen, J., M. Sato, R. Ruedy, K. Lo, D. W. Lea, and M. Medina-Elizade. 2006. Global temperature change. Proceedings of the National Academy of Sciences 103:14288–14293.

Helsel, D., and R. Hirsch. 2002. Statistical methods in water resources. Techniques of Water-Resources Investigations Book 4:395.

Herlihy, A. T., N. C. Kamman, J. C. Sifneos, D. Charles, M. D. Enache, and R. J. Stevenson. 2013. Using multiple approaches to develop nutrient criteria for lakes in the conterminous USA. Freshwater Science 32:367–384.

Hollister, J. W., D. Q. Kellogg, B. J. Kreakie, S. S. Shivers, B. W. Milstead, E. Herron, L. Green, and A. Gold. 2019. Zenodo - complete citation when paper submited.

Hollister, J. W., W. B. Milstead, and B. J. Kreakie. 2016. Modeling lake trophic state: A random forest approach. Ecosphere 7.

Hurlbert, S. H. 1984. Pseudoreplication and the design of ecological field experiments. Ecological monographs 54:187–211.

Jones, P. D., M. New, D. E. Parker, S. Martin, and I. G. Rigor. 1999. Surface air temperature and its changes over the past 150 years. Reviews of Geophysics 37:173–199.

Jones, P., and M. Hulme. 1996. Calculating regional climatic time series for temperature and precipitation: Methods and illustrations. International Journal of Climatology: A Journal of the Royal Meteorological Society 16:361–377.

Kellogg, D., A. J. Gold, S. Cox, K. Addy, and P. V. August. 2010. A geospatial approach for assessing denitrification sinks within lower-order catchments. Ecological Engineering 36:1596–1606.

Kosmala, M., A. Wiggins, A. Swanson, and B. Simmons. 2016. Assessing data quality in citizen science. Frontiers in Ecology and the Environment 14:551–560.

Kosten, S., V. L. Huszar, E. Bécares, L. S. Costa, E. Van Donk, L.-A. Hansson, E. Jeppesen, C. Kruk, G. Lacerot, N. Mazzeo, and others. 2012. Warmer climates boost cyanobacterial dominance in shallow lakes. Global Change Biology 18:118–126.

Kreeger, D. A., C. M. Gatenby, and P. W. Bergstrom. 2018. Restoration potential of several native species of bivalve molluscs for water quality improvement in mid-atlantic watersheds. Journal of Shellfish Research 37:1121–1158.

Litke, D. W. 1999. Review of phosphorus control measures in the united states and their effects on water quality. Water-Resources Investigations Report 99:4007.

Lottig, N. R., P.-N. Tan, T. Wagner, K. S. Cheruvelil, P. A. Soranno, E. H. Stanley, C. E. Scott, C. A. Stow, and S. Yuan. 2017. Macroscale patterns of synchrony identify complex relationships among spatial and temporal ecosystem drivers. Ecosphere 8:12.

Lottig, N. R., T. Wagner, E. N. Henry, K. S. Cheruvelil, K. E. Webster, J. A. Downing, and C. A. Stow. 2014. Long-term citizen-collected data reveal geographical patterns and temporal trends in lake water clarity. PLoS ONE 9:e95769.

Michalak, A. M., E. J. Anderson, D. Beletsky, S. Boland, N. S. Bosch, T. B. Bridgeman, J. D. Chaffin, K. Cho, R. Confesor, I. Daloğlu, and others. 2013. Record-setting algal bloom in lake erie caused by agricultural and meteorological trends consistent with expected future conditions. Proceedings of the National Academy of Sciences 110:6448–6452.

Nelson, K. C., and M. A. Palmer. 2007. Stream temperature surges under urbanization and climate change: Data, models, and responses 1. JAWRA Journal of the American Water Resources Association 43:440–452.

Nojavan, F., B. J. Kreakie, J. W. Hollister, and S. S. Qian. 2019. Rethinking the lake trophic state index. PeerJ Preprints.

Oliver, S. K., S. M. Collins, P. A. Soranno, T. Wagner, E. H. Stanley, J. R. Jones, C. A. Stow, and N. R. Lottig. 2017. Unexpected stasis in a changing world: Lake nutrient and chlorophyll trends since 1990. Global Change Biology 23:5455–5467.

Paerl, H. W., and J. Huisman. 2009. Climate change: A catalyst for global expansion of harmful cyanobacterial blooms. Environmental Microbiology Reports 1:27–37.

Paerl, H. W., J. T. Scott, M. J. McCarthy, S. E. Newell, W. S. Gardner, K. E. Havens, D. K. Hoffman, S. W. Wilhelm, and W. A. Wurtsbaugh. 2016. It takes two to tango: When and where dual nutrient (N & P) reductions are needed to protect lakes and downstream ecosystems. Environmental Science & Technology 50:10805–10813.

Pennino, M. J., R. I. McDonald, and P. R. Jaffe. 2016. Watershed-scale impacts of stormwater green infrastructure on hydrology, nutrient fluxes, and combined sewer overflows in the mid-atlantic region. Science of the Total Environment 565:1044–1053.

Read, E. K., V. P. Patil, S. K. Oliver, A. L. Hetherington, J. A. Brentrup, J. A. Zwart, K. M. Winters, J. R. Corman, E. R. Nodine, R. I. Woolway, and others. 2015. The importance of lake-specific characteristics for water quality across the continental United States. Ecological Applications 25:943–955.

Reisinger, A. J., E. Woytowitz, E. Majcher, E. J. Rosi, K. T. Belt, J. M. Duncan, S. S. Kaushal, and P. M. Groffman. 2019. Changes in long-term water quality of baltimore streams are associated with both gray and green infrastructure. Limnology and Oceanography 64:S60–S76.

Rhode Island State Legislature. 1995. Phosphate reduction act of 1995.

Schindler, D. 2009. Lakes as sentinels and integrators for the effects of climate change on watersheds, airsheds, and landscapes. Limnology and Oceanography 54:2349–2358.

Schindler, D. W., R. Hecky, D. Findlay, M. Stainton, B. Parker, M. Paterson, K. Beaty, M. Lyng, and S. Kasian. 2008. Eutrophication of lakes cannot be controlled by reducing nitrogen input: Results of a 37-year whole-ecosystem experiment. Proceedings of the National Academy of Sciences 105:11254–11258.

Smith, V. H. 2003. Eutrophication of freshwater and coastal marine ecosystems a global problem. Environmental Science and Pollution Research 10:126–139.

Soranno, P. A., L. C. Bacon, M. Beauchene, K. E. Bednar, E. G. Bissell, and al. et. 2017. LAGOS-NE: A multi-scaled geospatial and temporal database of lake ecological context and water quality for thousands of US lakes. Gigascience 6.

Soranno, P. A., E. G. Bissell, K. S. Cheruvelil, S. T. Christel, S. M. Collins, C. E. Fergus, C. T. Filstrup, J.-F. Lapierre, N. R. Lottig, S. K. Oliver, and others. 2015. Building a multi-scaled geospatial temporal ecology database from disparate data sources: Fostering open science and data reuse. GigaScience 4:28.

Stachelek, J., and S. Oliver. 2017. LAGOSNE: Interface to the lake multi-scaled geospatial and temporal database, R package version 1.1.0. https://cran.r-project.org/package=LAGOSNE.

Stoddard, J. L., J. Van Sickle, A. T. Herlihy, J. Brahney, S. Paulsen, D. V. Peck, R. Mitchell, and A. I. Pollard. 2016. Continental-scale increase in lake and stream phosphorus: Are oligotrophic systems disappearing in the united states? Environmental Science & Technology 50:3409–3415.

Taranu, Z. E., I. Gregory-Eaves, P. R. Leavitt, L. Bunting, T. Buchaca, J. Catalan, I. Domaizon, P. Guilizzoni, A. Lami, S. McGowan, and others. 2015. Acceleration of cyanobacterial dominance in north temperate-subarctic lakes during the anthropocene. Ecology Letters 18:375–384.

Vitousek, P. M., J. D. Aber, R. W. Howarth, G. E. Likens, P. A. Matson, D. W. Schindler, W. H. Schlesinger, and D. G. Tilman. 1997. Human alteration of the global nitrogen cycle: Sources and consequences. Ecological Applications 7:737–750.

Wasserstein, R. L., N. A. Lazar, and others. 2016. The ASA’s statement on p-values: Context, process, and purpose. The American Statistician 70:129–133.

Williamson, C. E., W. Dodds, T. K. Kratz, and M. A. Palmer. 2008. Lakes and streams as sentinels of environmental change in terrestrial and atmospheric processes. Frontiers in Ecology and the Environment 6:247–254.

Yang, Y.-Y., and M. G. Lusk. 2018. Nutrients in urban stormwater runoff: Current state of the science and potential mitigation options. Current Pollution Reports 4:112–127.

Yuan, L. L., A. I. Pollard, S. Pather, J. L. Oliver, and L. D’Anglada. 2014. Managing microcystin: Identifying national-scale thresholds for total nitrogen and chlorophyll a. Freshwater biology 59:1970–1981.

